# Characterization of a Novel Pseudomonad with Biocontrol Activity Against *Aphanomyces euteiches*

**DOI:** 10.64898/2026.05.18.726007

**Authors:** Ashlyn Kirk, Sean D. Workman, Ava M. Tiefenbach, Sean M. Hemmingsen, Christopher K. Yost

## Abstract

*Aphanomyces euteiches*, the causative agent of *Aphanomyces* root rot (ARR), is of major concern for pea and other legume crops globally. This oomycete pathogen causes substantial decreases in crop yields, is unaffected by most fungicides, and persists in the soil for many years via its resilient oospores. Given the significance of pea crops in sustainable agriculture, namely the ability to fix nitrogen and act as a sustainable protein source, solutions to ARR are of high importance. We used RNA-seq in a novel strain of *Pseudomonas donghuensis* to identify two biosynthetic gene clusters under GacA/S control that are involved in producing bioactive molecules capable of inhibiting *A. euteiches*. Based on similarity to other reported clusters in *Pseudomonas*, the first is predicted to encode for a pseudoiodinine compound, while the second is predicted to produce the siderophore 7-hydroxytropolone. Individual knockouts of each cluster showed loss of inhibitory action of *P. donghuensis* NRC29 against *A, euteiches in vivo*. This is the first report highlighting the potential of *P. donghuensis* and the products of the two identified biosynthetic pathways as biocontrol agents for *A. euteiches*. Further investigations into the efficacy of *P. donghuensis* NRC29 and its metabolites in inhibiting *A. euteiches* in field trials will be of high value in developing sustainable strategies for ARR mitigation.

**Importance:** Modern fungicidal treatments for control of root rot in pulse crops are ineffective for control of *A. euteiches*, leaving limited strategies for management of *A. euteiches* infected fields. We describe a novel *P. donghuensis* strain with potential for biocontrol against this persistent pathogen. Given the economic value of peas and other pulses globally, further work into harnessing the bioactive metabolites produced by this strain into a practical in-field treatment will be valuable.

## Introduction

*Aphanomyces euteiches* is a soilborne oomycete pathogen that infects plant roots in the family Fabaceae, including field peas (*Pisum sativum*), lentils (*Lens culinaris*), common beans (*Phaseolus vulgaris*), chickpeas (*Cicer arietinum*) and other economically important pulse crops (1, 2). Infection of host plants by *A. euteiches* leads to *Aphanomyces* root rot (ARR), a disease that typically results in stunted seedlings, yellowing of leaves, and soft water-soaked lateral roots that display honey-brown discolouration (3, 4). *A. euteiches* is one of the most destructive pea pathogens and ARR is one of the major limitations to global pea production. The symptoms of ARR are often compounded through the coinfection of pea roots by fungal pathogens such as *Fusarium* spp. and is commonly referred to as a root rot complex (1). Crop yields can be reduced by over 80% when root rot is particularly severe (5, 6). Besides their economic importance, with an estimated 14.5 million tonnes produced by global leaders in pea production in 2025 (7), peas and other legume crops also play a vital role in sustainable agriculture. Peas form important interactions with bacteria from the genus *Rhizobium*, leading to the formation of root nodules that allow for fixation of nitrogen from the atmosphere. These interactions reduce the need for chemical fertilizers and decrease the risk of eutrophication by inorganic nitrogen fertilizers (8), making mitigation of diseases of multifaceted importance.

Despite being recognized as a serious disease of pea since the 1920s (9), and having been reported in all the major pea cultivation regions of the world, no broadly effective method has been developed for the management of ARR. Management of ARR is made difficult by the resilient nature of *A. euteiches* oospores, which are able to survive during harsh environmental conditions and can remain viable in the soil for over 10 years (4, 5). Additionally, given key differences between fungi and oomycetes, including lack of commonly targeted fungal membrane sterols in oomycetes and cell walls comprised of cellulose rather than chitin, most agricultural fungicides designed to target fungal pathogens are ineffective against oomycetes. Historically, the limited strategies available to producers have been avoidance of *A. euteiches* infected fields or long crop rotations with non-host crops such as wheat or oats to reduce inoculum abundance (1). Efforts to breed resistant cultivars of pea are ongoing, and though pea germplasms with partial resistance have been identified (10–12), progress is limited. A handful of foliar fungicides and fungicidal seed treatments such as INTEGO^®^ Solo, Vibrance^®^ Maxx with INTEGO^®^, Zeltera^®^ Pulse, and Rancona Trio have been approved for the early season suppression of ARR in Canada (2, 13), but their inconsistent and limited control of the disease has led to an interest in biological control agents as a potential alternative (14, 15).

Biological control agents aim to harness the rich chemical diversity of the microbial world to suppress plant pathogens while minimizing negative effects on the beneficial members of the rhizosphere. The development of bacterial or fungal isolates as seed coatings or soil inoculants has led to successful commercial products such Integral (*Bacillus amyloliquefaciens* MBI 600) showing effectiveness against *Rhizoctonia* spp. and *Fusarium* spp. (16), and Xedavir (*Trichoderma asperellum*) repressing *Pythium* spp., *Phitophtroa capsica*, and other causative agents of root rot (17). Biological control agents effective against *A. euteiches* are of major importance, especially with the increasing demand to find sustainable protein alternatives such as pea protein (18). Several bacterial strains antagonistic to *A. euteiches* have been identified and a recent meta-analysis determined that while there is significant heterogeneity in their effectiveness in controlling ARR, there is evidence that biological control methods hold promise for the management of ARR overall (14).

In this work, we characterize a novel strain of *Pseudomonas donghuensis*, isolated from Canadian prairie soil with a history of pea crop cultivation, that shows significant inhibition of *A. euteiches*. We use CRISPR-RNA guided integrases to generate targeted knockouts of a two-component system previously implicated in the biosynthesis of antifungal secondary metabolites, and take a multiomics approach to identify the genomic loci implicated in the production of two antioomycete compounds.

## Results & Discussion

### The Gac two-component system controls production of antioomycete compounds

The Gac two-component system is a global regulatory system found conserved across Gram-negative bacteria and consists of a response regulator (GacA) and sensor kinase (GacS) that work in tandem to regulate a variety of physiological responses to environmental stimuli (19). Processes involve regulation of N-acyl homoserine lactone and secondary metabolite production, motility, biofilm formation, and regulation of type 3 and type 6 secretion systems (19). Based on previous accounts of high-level regulation by GacA/S of anti-fungal compounds described in pseudomonads (20), we targeted the Gac two-component system to determine if it also regulated antioomycete activity in *P. donghuensis* NRC29. Knockout of *gacA* in *P. donghuensis* NRC29 resulted in loss of anti-mycelial inhibition of *A. euteiches* on potato dextrose agar (PDA) in comparison to the wildtype (Figure 1). Complementation of *gacA* carried on pSEVA231 resulted in restoration of the antioomycete phenotype, confirming regulation of antioomycete compounds by the GacS/A signal cascade (Figure 1). Knockout and complementation of *gacS* yielded similar results (data not shown).

**Figure 1.**
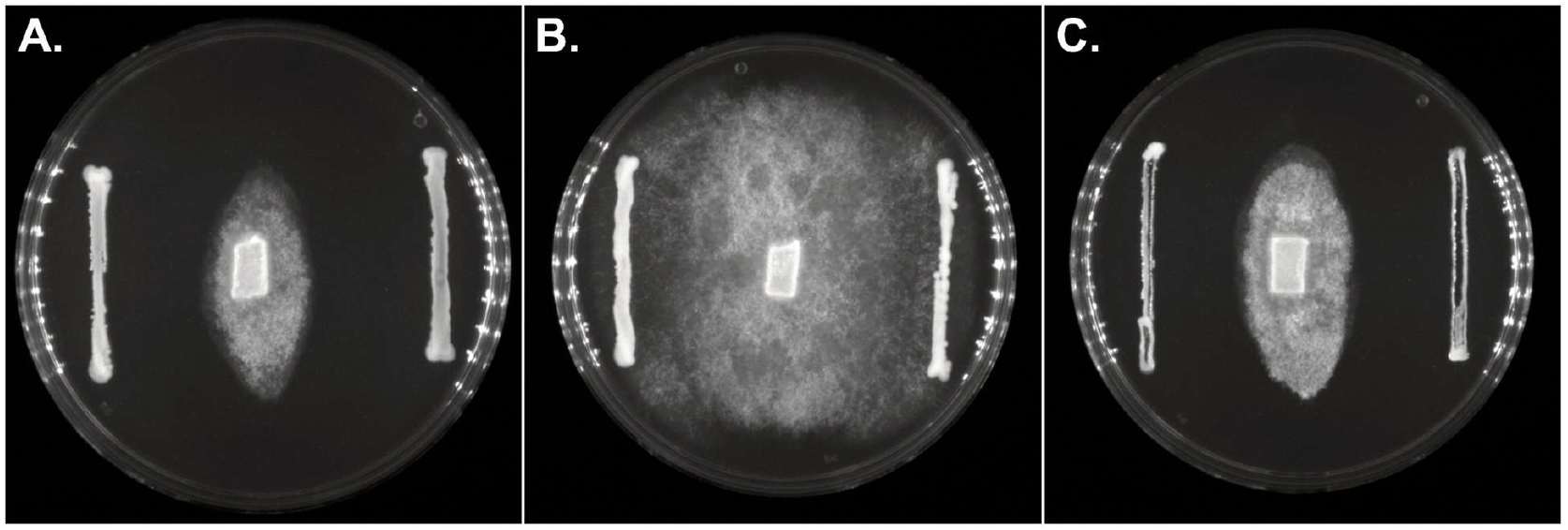
Differential inhibition of *A. euteiches* mycelial growth on PDA by *P. donghuensis* NRC29 strains. **A.** Wildtype *P. donghuensis* NRC29 **B**. Δ*gacA P. donghuensis* NRC29 **C**. Δ*gacA P. donghuensis* NRC29 with *gacA* provided in trans on pSEVA231

Inhibition assays using spent media fractions were done to help elucidate the chemical nature of the bioactive compound(s). Wildtype *P. donghuensis* NRC29 showed strongest activity in the 50% methanol step elution (Supplementary Figure 1), suggesting that the inhibitory activity is associated with one or more small molecules, rather than proteins or cyclic peptides. As with our culture-based assays, inhibition was lost in all fractions taken from the Δ*gacA* mutant and restored when *gacA* was provided *in trans* (Supplementary Figures 2 & 3).

### RNA-Seq uncovers differentially regulated biosynthetic gene clusters in the Δgac mutants

Disruption of antioomycete capabilities in Δ*gacA P. donghuensis* NRC29 prompted investigation into the expression of downstream metabolic pathways in the absence of GacA activity that could lead to loss of activity. RNAseq data was collected from both the log and stationary phases of wild type NRC29 and Δ*gacA* NRC29 grown in potato dextrose broth (PDB) supplemented with M9 salts. Comparison of the 75 most variably expressed genes across the four datasets show minimal transcriptional differences in log-phase of growth between wildtype NRC29 and Δ*gacA* NRC29 (Figure 2), however, significant differences can be seen during stationary phase (Figure 2). Given the role of GacA/S in regulating secondary metabolic processes, differences in downstream regulation during stationary phase are expected in comparison to log-phase as cells transition from a primarily focused growth physiology to a survival and competition-centered metabolism (21). Previous studies have demonstrated the tight regulation of *gacA* expression in tandem with bacterial growth, with highest expression levels of *gacA* observed during stationary phase (19, 22, 23).

**Figure 2.**
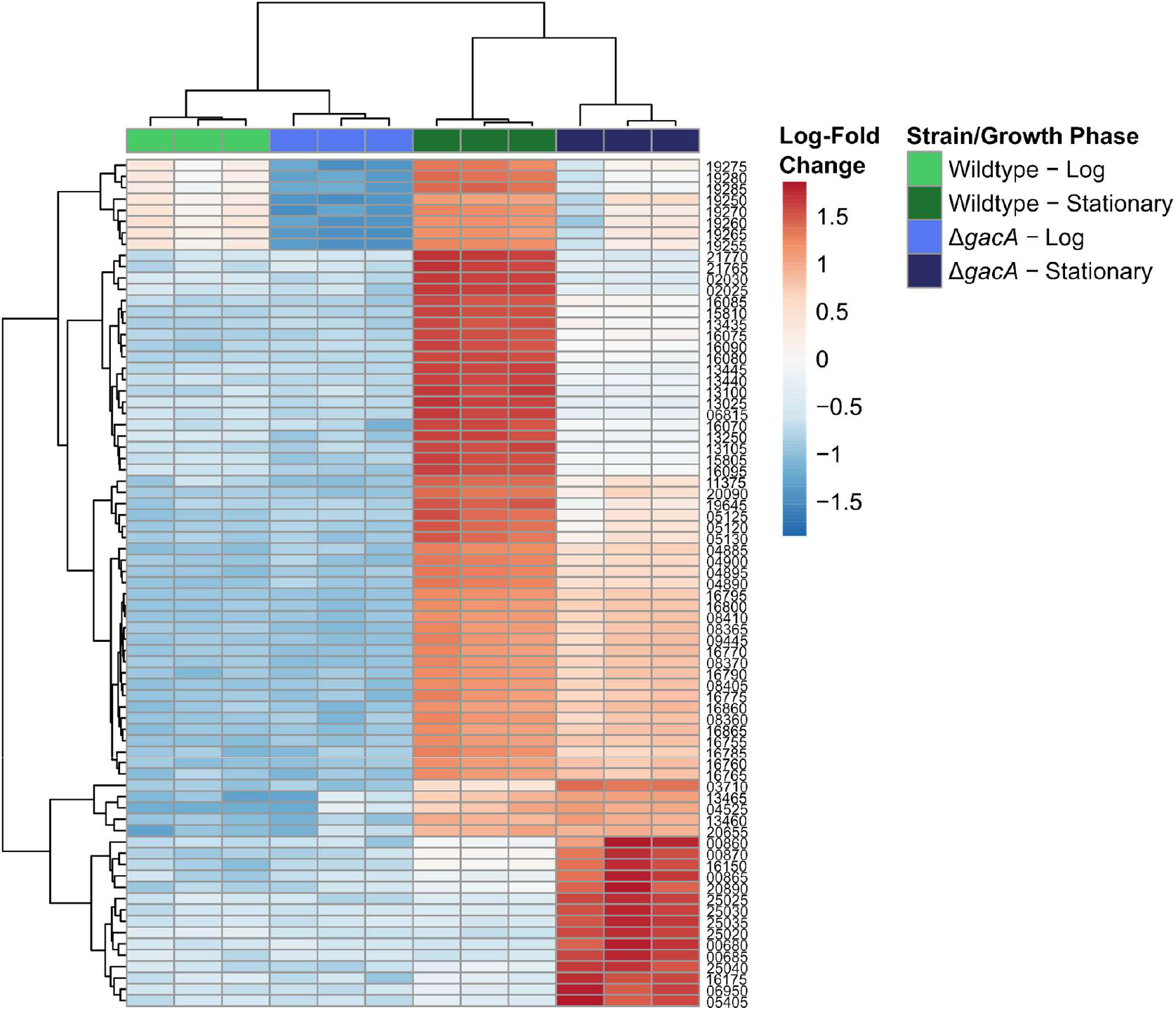
RNA-Seq heatmap showing top 75 most variably expressed genes across all datasets. Hierarchical clustering reveals enhanced transcriptional differences between wildtype and Δ*gacA* strains during stationary phase.

Among differentially expressed loci during stationary phase, we identified two biosynthetic gene clusters (BGCs) of interest in which expression was substantially reduced in the NRC29 Δ*gacA* mutant strain (Figure 3A, 3B). Both BGCs have been previously linked to GacA/S control and synthesize products previously identified to have biocontrol activity against some bacterial and fungal plant pathogens (20, 24, 25). A search of both clusters against NCBI’s core_nt database using default parameters show that both are restricted to *Pseudomonas* (Supplementary Tables 1 & 2).

**Figure 3.**
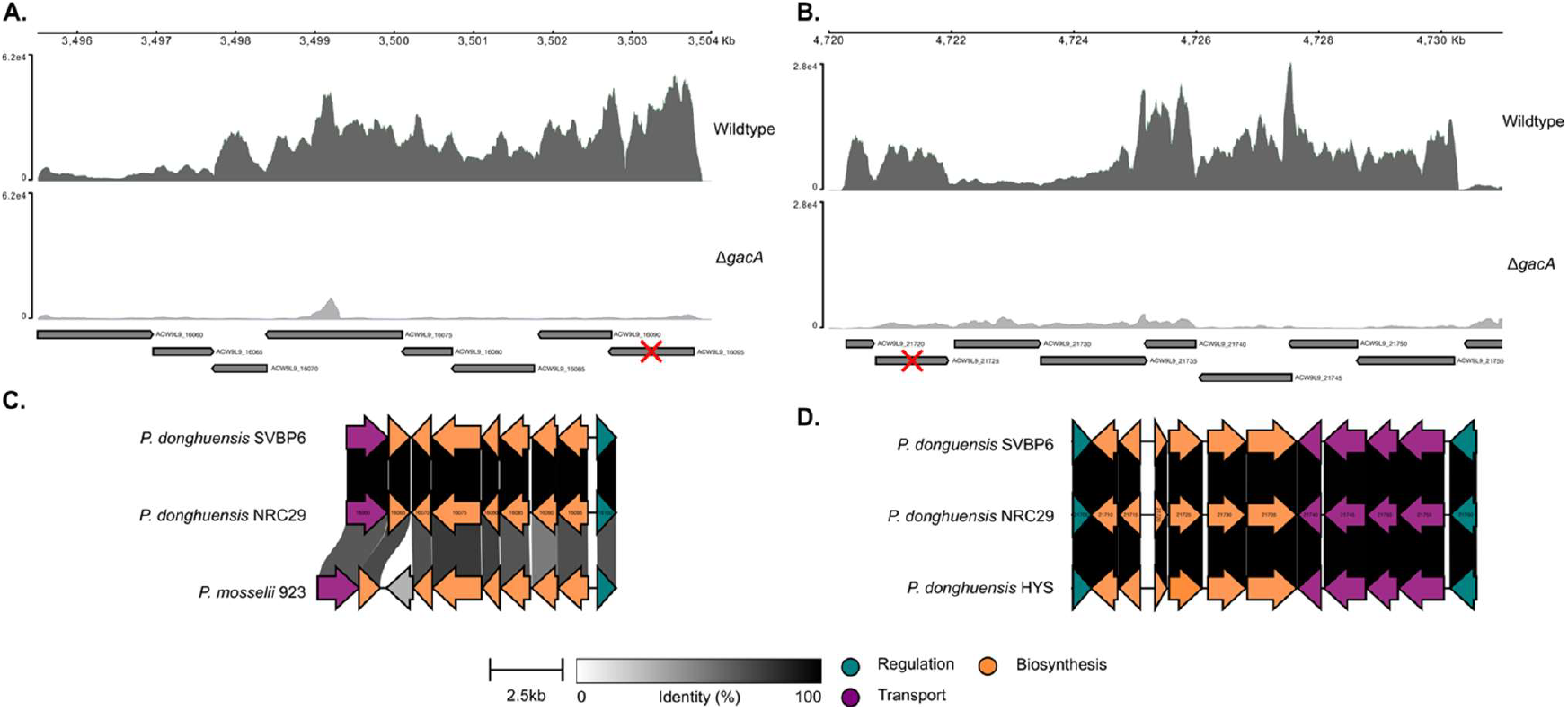
Visualization of RNA-Seq count data at genetic loci of interest and comparison of biosynthetic gene clusters (BGCs) found in *P. donghuensis* NRC29 to homologs. **A.** ACW9L9_16095 encodes a putative tetracenomycin polyketide synthesis gene. **B**. ACW9L9_21725 encodes an acyl-CoA dehydrogenase previously implicated in the biosynthesis of siderophores. Red Xs indicate the loci targeted for insertional mutagenesis. **C**. Comparison of pseudoiodinine BGC in NRC29 to previously identified homologs. **D**. Comparison of 7-hydroxytropolone BGC in NRC29 to previously identified homologs.

The first BGC, shares 99.67% identity (100% coverage) with a 9-gene BGC identified in *P. donghuensis* SVBP6 (20) (Figure 3C) known to direct the biosynthesis, regulation, and transport of a pseudoiodinine compound (24, 26). *P. donghuensis* SVBP6 shows activity against several fungal plant pathogens including *Fusarium, Macrophomina, Colletotrichum, Phomopsis*, and *Cercospora* (27). Additionally, both culture and supernatant of the pseudoiodinine producing *P. mosselii* 923 has shown efficacy in inhibiting *Xanthomonas spp*. and *Magnaporthe oryzae in planta* in both greenhouse and field trials in rice (24). Activity of purified pseudoiodinine has not yet been reported, making this compound of interest for further isolation and testing against *A. euteiches* and other root-rot causing pathogens.

The second BGC of interest is predicted to encode the siderophore 7-hydroxytropolone (7-HT) based on similarity to previously described BGCs in *P. donghuensis* (20, 28–30) (Figure 3D). Biocontrol studies of 7-HT have centred primarily on the antibacterial activity against *Dickeya solani*, the causative agent of blackleg disease in potatoes (28, 31, 32) as well as select fungal phytopathogens including *Fusarium* and *Macrophomina* (33). Like pseudoiodinine, the inhibitory effects of 7-HT have not yet been tested against *A. euteiches*, making further testing of the isolated compound of significant interest.

### Abundant iron access masks the effects of antioomycete compounds

To further clarify the metabolic pathway contributing to inhibition of *A. euteiches* by *P. donghuensis* NRC29, we created knockouts of the two previously identified BGCs by creating CRISPR guided transposon disruptions in two differentially expressed loci identified in our RNA-seq assays (Figure 3). ACW9L9_16095, part of the putative pseudoiodinine BGC is predicted to encode a tetracenomycin polyketide synthesis 8-O-methyltransferase, while ACW9L9_21725, found in the predicted 7-HT siderophore cluster, is predicted to encode for an acyl-coA dehydrogenase (Figure 3). Since siderophore biosynthesis is heavily repressed in the presence of ferrous ion via the ferric uptake regulator (Fur) pathway in pseudomonads (34), inhibition assays of wildtype and knockout strains were carried out on both 0.5X PDA as well as 0.5X PDA supplemented with FeCl_3_ to better resolve which molecules contribute to *A. euteiches* inhibition.

To determine the appropriate FeCl_3_ concentration required to inhibit siderophore production, we first tested a gradient of concentrations in *A. euteiches* inhibition assays with wildtype NRC29 (Figure 4). Concentration-dependent loss of inhibition against *A. euteiches* was observed, with the greatest loss of inhibition seen at 50 μM (Figure 4). A concentration of 50 μM FeCl_3_ was, therefore, used for the rest of our assays to compare inhibition of the two knockout strains under iron-rich and iron-limited conditions.

**Figure 4.**
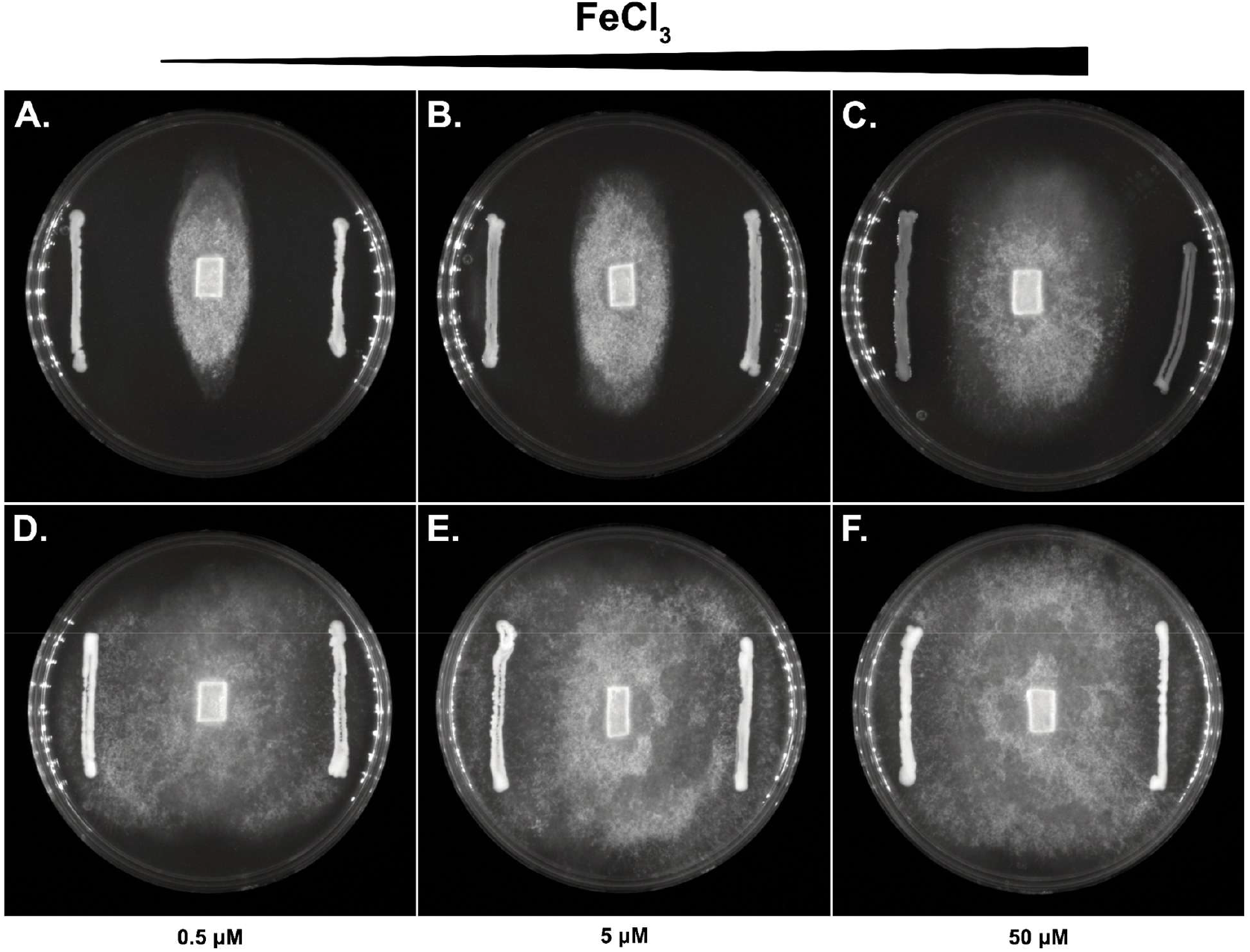
Iron scavenging siderophores contribute significantly to *A. euteiches* inhibition by *P. donghuensis* NRC29. **A-C.** *A. euteiches* is more resilient to inhibition by *P. donghuensis* NRC29 as iron concentration increases. **D-F**. *A. euteiches* growth is not impacted by Δ*gacA P. donghuensis* NRC29 at all iron concentrations.

For ΔACW9L9_21725, inhibition levels between iron-rich and iron-deficient media remained similar, which is consistent with our hypothesis that a non-iron-dependent molecule such as pseudoiodinine is involved in inhibition (Figure 5). In contrast, ΔACW9L9_16095, which was constructed to eliminate expression of the pseudoiodinine cluster but retain 7-HT expression, showed active inhibition without iron, but almost a total loss of inhibition in the presence of iron (Figure 5). Previous studies examining siderophore production via absorbance of culture supernatants of *P. donghuensis* have shown approximately 8-10 μM ferrous ions are required to suppress 7-HT production to minimum levels in King’s B medium, while only 1-2 μM are needed to suppress pyoverdine production to the same degree (25, 30). Collectively, we can infer that 7-HT production is contributing to *A. euteiches* inhibition by *P. donghuensis* NRC29 and expression is minimal in the iron supplemented condition.

**Figure 5.**
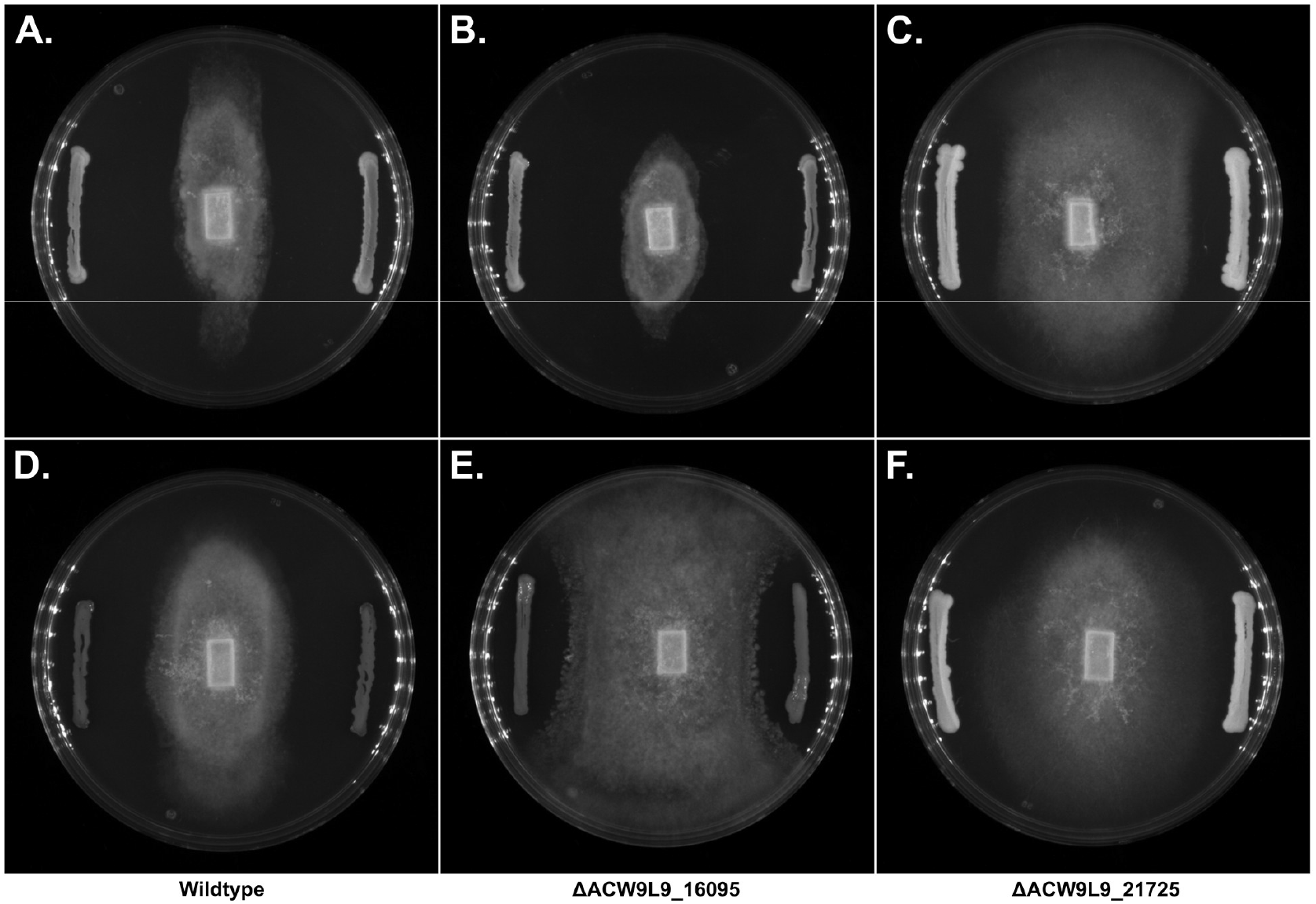
Differential phenotypes of *P. donghuensis* NRC29 insertional knockouts. **A-C.** Wildtype, ΔACW9L9_16095, and ΔACW9L9_21725 *P. donghuensis* plated on 0.5x PDA. **D-F**. Wildtype, ΔACW9L9_16095, and ΔACW9L9_21725 *P. donghuensis* plated on 0.5x PDA supplemented with 50 μM FeCl3.

Since 7-HT production is inhibited by abundant iron, future field trials will need to include considerations of the limitation on siderophore production from culture inoculants in iron-rich soils. Efficacy trials of purified 7-HT and pseudoiodinine will be a valuable next step to determine if these compounds are bioactive independent of other cellular processes or in iron-abundant conditions. Use of purified compounds could also avoid potential environmental limitations on the GacA/S signal cascade, or loss of function mutations during inoculum preparation (35).

Notably, we still observed a modest level of inhibition in the iron-rich condition of our ΔACW9L9_16095 assay (Figure 5), but not in the GacA knockouts (Figure 4). The GacA/S signal cascade regulates a wide variety of secondary metabolites in pseudomonads (36), so, it is likely that there are additional active molecules under GacA control contributing to *Aphanomyces* inhibition in addition to 7-HT and pseudoiodinine. AntiSMASH (37) of the *P. donghuensis* NRC29 genome using the default settings found a high-identity match to a hydrogen cyanide biosynthetic cluster found broadly distributed across pseudomonads. This cluster is under GacA/S control (38) and has been implicated in mediation of biocontrol in *Pseudomonas* (39– 41), making it a plausible contributor to *A. euteiches* inhibition by *P. donghuensis* NRC29 in the absence of 7-HT and pseudoiodinine activity.

## Conclusion

Microbial natural products offer an environmentally sustainable alternative to traditional chemical biocide treatments and help protect beneficial plant-associated organisms. This study demonstrates the potential of natural products produced by *P. donghuensis* NRC29 for use in biocontrol of *A. euteiches*, specifically, 7-HT and pseudoiodinine. Given the magnitude and longevity of destruction caused by *A. euteiches* on peas and other pulse crops and lack of available treatments, further work into development of *P. donghuensis* NRC29 or its metabolites for field-deployable biocontrol is of high value.

## Materials & Methods

### Bacterial strains and culture conditions

Biocontrol strain *Pseudomonas donghuensis* NRC29, was cultured from a soil sample taken from a prairie research farm with extensive pea crop cultivation. The wildtype and its derivative mutants were propagated at 30°C in LB broth or on 1.5% LB agar. *Escherichia coli* strains were grown at 30°C or 37°C in LB broth or on 1.5% LB agar. *E. coli* DH5α was used for the construction and propagation of both mutagenesis and complementation plasmids, *E. coli* MFD*pir* was used for the conjugation of plasmids into *P. donghuensis* NRC29, and *E. coli* K12 MG1655 was used as a negative control in *A. euteiches* inhibition assays. Media for *E. coli* MFD*pir* were supplemented with 0.3 mM 2,6-diaminopimelic acid (DAP). When required, growth media were supplemented with 50 μg/mL apramycin, 50 μg/mL kanamycin, 100 μg/mL spectinomycin, or 5 μg/mL triclosan.

### Construction of mutagenesis and complementation plasmids

Insertional knockouts were generated using RNA-guided CRISPR-transposons (42). Spacer sequences targeting genes of interest were designed using the CAST Guide RNA Tool (42) and oligonucleotides were designed to preserve the functional CRISPR array after BsaI digestion. Forward spacer sequences were prepended with 5’-ATAAC-3’ and had a single G appended at the 3’ end. The reverse complementary spacer sequences were prepended with 5’-TTCAC-3’ and had a single G appended at the 3’end. Complementary spacer oligonucleotides were mixed at 20 μM in 10 mM Tris-HCl (pH 9.0), 0.1 mM EDTA, 50 mM NaCl and heated at 95°C for 5 minutes before cooling to 4°C at a rate of 0.1°C/s. Duplex DNA was quantified using a Qubit dsDNA Broad Range Assay Kit and duplex spacers were then cloned into a pSL1521 derivative delivering an apramycin resistance using Golden Gate cloning with BsaI-HFv2 and T4 DNA ligase (NEB). Following the completion of the Golden Gate cloning protocol, 1 μL of fresh BsaI-HFv2 was added to each reaction and allowed to digest at 37°C for 1 hour before heat inactivation at 80°C for 20 minutes. Heat inactivated reactions were transformed into *E. coli* DH5α and colonies were screened for successful incorporation of spacer sequences using colony PCR. Confirmed colonies were grown overnight at 30°C in LB media supplemented with apramycin and spectinomycin for plasmid purification. Complementation plasmids for Δ*gacA* and Δ*gacS* mutants were constructed using a pSEVA231 backbone (43). The ORFs encoding *gacA* and *gacS* along with 500 bp both upstream and downstream of the annotated start and end sites of the ORFs were PCR amplified from purified NRC29 genomic DNA using Q5 High-Fidelity Polymerase (NEB) and primers that incorporated restriction cleavage sites for EcoRI and HindIII. PCR products were purified using a QIAquick PCR Purification Kit (QIAGEN). Purified PCR products and pSEVA231 were each double digested using EcoRI-HF and HindIII-HF (NEB) at 37°C for 1 hour before heat inactivation at 80°C for 20 minutes. Digested PCR products were ligated into digested pSEVA231 overnight using T4 DNA ligase (NEB) at 16°C. Ligation reactions were transformed into *E. coli* DH5α and individual colonies were picked and grown overnight at 37°C in LB media supplemented with kanamycin for plasmid purification. All plasmid sequences were confirmed using whole plasmid sequencing performed by Plasmidsaurus.

### Chemical fractionation

Cell-free aliquots of spent media from wildtype, Δ*gacA*, and *gacA* complemented Δ*gacA P. donghuensis* NRC29 cultures were lyophilized and subjected to crude chemical fractionation. The lyophilized media was resuspended in water and the soluble fraction from a subsequent 80% ethanol precipitation was loaded on to a C18 column before being subjected to 0%, 50% and 100% methanol step elutions.

### Aphanomyces euteiches mycelial growth inhibition assays

*A. euteiches* inhibition assays were carried out on 0.5x PDA. The plates were inoculated with *A. euteiches* by placing an agar plug from a 0.5x PDA plate containing mycelial growth at the center of the test plate. Individual colonies of *P. donghuensis* NRC29, its derivative mutants, or *E. coli* K12 MG1655 were streaked at the edges of the 0.5x PDA plate and the plates were allowed to incubate at 23°C in a dark incubator for 5 days before measurement of inhibition. For inhibition assays using spent media, lyophilized material from fractionated spent media was solubilized using filter sterilized DMSO and 50 μL of each solubilized fraction was dispensed on to a sterile paper disks placed on a 0.5x PDA inoculated with *A. euteiches* as above.

### Insertional mutagenesis of NRC29

CAST plasmids were transformed into *E. coli* MFD*pir* and individual colonies were picked and grown overnight at in LB media for conjugation. 1 mL aliquots of overnight cultures of *E. coli* MFD*pir* harboring CAST plasmids and of NRC29 were separately centrifuged for 3 minutes at 8,000 RCF and the supernatant was removed. Cell pellets were resuspended in 1 mL sterile phosphate buffered saline (PBS) and centrifuged for 3 minutes at 8,000 RCF again. The conjugation was initiated by resuspending the *E. coli* MFD*pir* cell pellet in 100 μL sterile PBS and using the cell suspension to resuspend the NRC29 cell pellet. The conjugation mixture was spotted on to LB agar supplemented with DAP and allowed to proceed at 30°C for 5 hours. The conjugation spots were then scraped and resuspended in 1 mL sterile PBS and 50 μL of the mixture was used to inoculate LB supplemented with triclosan and apramycin. After 24 hours of growth at 30°C, the cultures were streaked on to LB agar supplemented with triclosan and apramycin. Colony PCR was used to screen individual colonies for successful insertional mutagenesis. PCR confirmed colonies were grown overnight in LB media supplemented with triclosan and apramycin to create glycerol stocks for storage of mutant strains at -80°C.

### RNA-seq

Wildtype NRC29, Δ*gacA* NRC29, and Δ*gacS* NRC29 were grown overnight in LB media and were used to inoculate 50 mL 0.5x potato dextrose broth (PDB) supplemented with 0.5x M9 salts to a starting optical density (OD) of 0.025 in triplicate. Cultures were grown at 30°C with 225 rpm shaking. When cultures had reached either log or stationary phase, 1 mL of each culture was added to 2 mL of RNAprotect Bacteria Reagent (QIAGEN) for stabilization. Total RNA was purified using an RNeasy Mini Kit (QIAGEN) following the manufacturer’s instructions. Ribosomal RNA was depleted using a NEBNext rRNA Depletion Kit (NEB), sequencing libraries were constructed using NEBNext Ultra II Directional RNA Library Prep Kit for Illumina (NEB) and were indexed with unique dual indices using NEBNext Multiplex Oligos for Illumina (NEB). Constructed libraries were checked for adapter contamination and insert size using an Agilent 2100 Bioanalyzer with a high sensitivity DNA chip and were qPCR quantified using the NEBNext Library Quant Kit for Illumina (NEB). Pooled libraries were shipped to the Michael Smith Genome Sciences Centre in Vancouver, British Columbia and sequenced on a NovaSeq platform (Illumina) targeting 150 million 150 base-pair paired-end reads for the combined pool. Sequencing reads were quality checked using FastQC and were aligned to the NRC29 genome using bwa-mem. The number of read pairs unambiguously mapping to each gene was determined using featureCounts. DESeq2 was used to normalize raw counts and test for differential expression of genes between wildtype and Δ*gac*A/Δ*gacS* mutants. A log-fold change threshold for individual Wald tests was set to 1.0 and a false detection rate cutoff of 0.1 was used to determine statistical significance.

## Acknowledgements

Funding was provided through the National Research Council of Canada Special Projects Program (SPP-117; Mechanisms of *Aphanomyces euteiches* biocontrol in diverse bacterial strains) and through an NSERC Discovery Grant to CKY. We greatly appreciate the contribution of Dr. A. Pagé for provision of bacterial strains.

## Data Availability

*P. donghuensis* NRC29 wgs reads are available through NCBI’s Sequence Read Archive under BioProject accession PRJNA1378988. RNA-seq datasets generated during this study are available in NCBI’s Gene Expression Omnibus under series GSE316785.

## Supplementary Material

**Supplementary Figure 1.**
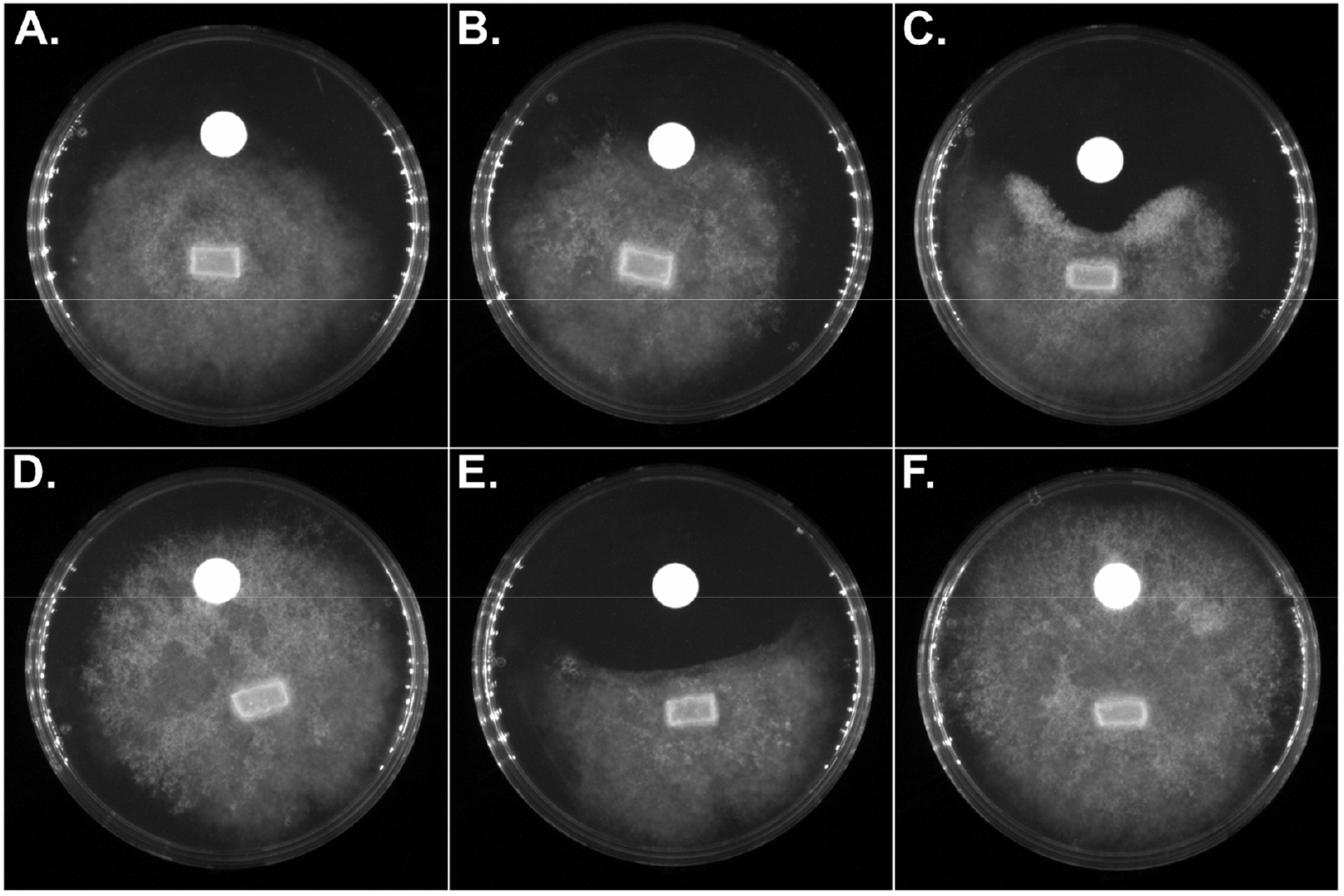
Inhibition assays of fractionation layers from wildtype *P. donghuensis* NRC29 against *A. euteiches*. **A**. Freeze-dried culture. **B**. 80% EtOH precipitation fraction. **C**. 80% EtOH soluble fraction. **D**. Water elution from C18 SPE column. **E**. 50% methanol elution from C18 SPE column. **F**. 100% methanol elution from C18 SPE column.

**Supplementary Figure 2.**
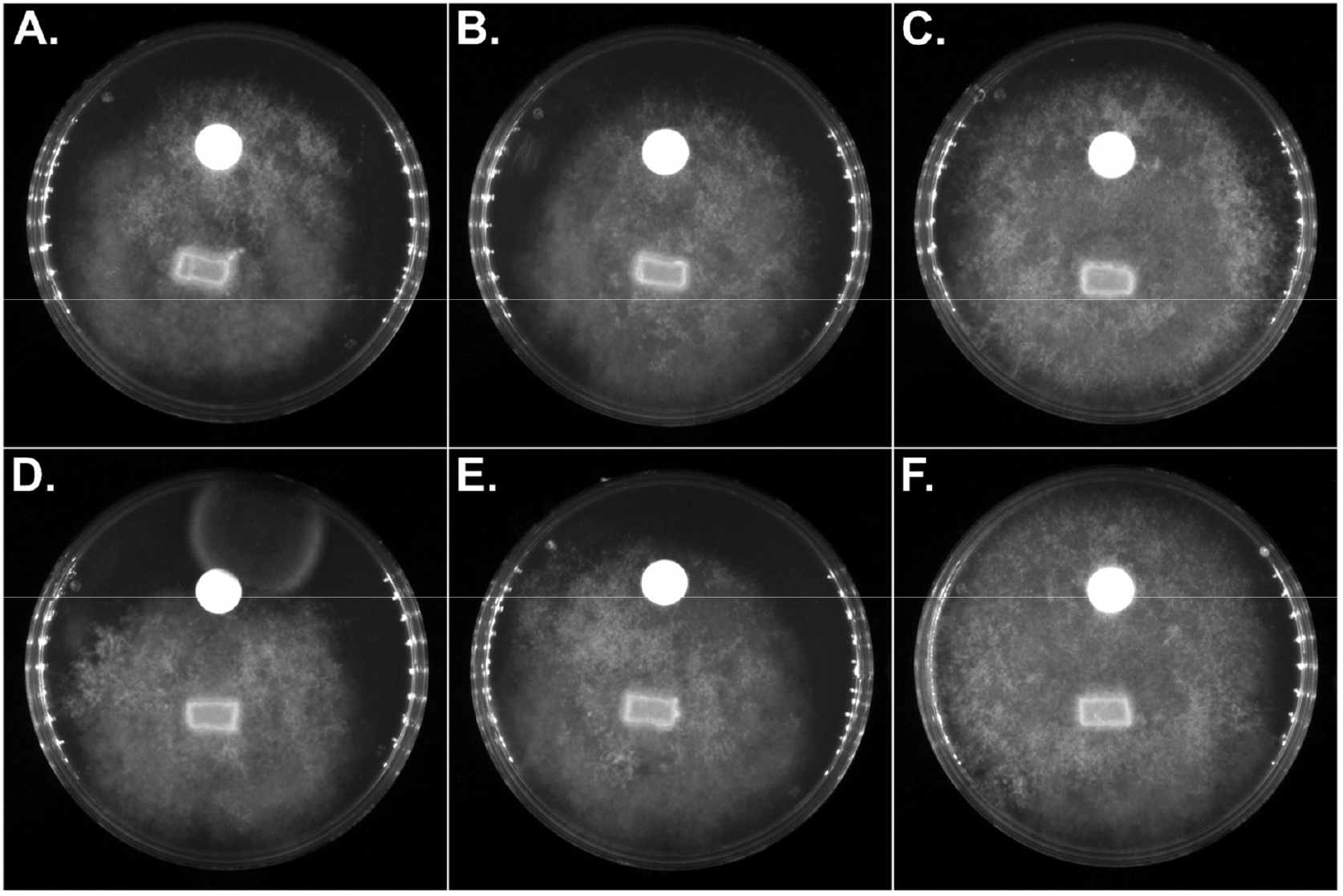
Inhibition assays of fractionation layers from Δ*gacA P. donghuensis* NRC29 against *A. euteiches*. **A**. Freeze-dried culture. **B**. 80% EtOH precipitation fraction. **C**. 80% EtOH soluble fraction. **D**. Water elution from C18 SPE column. **E**. 50% methanol elution from C18 SPE column. **F**. 100% methanol elution from C18 SPE column.

**Supplementary Figure 3.**
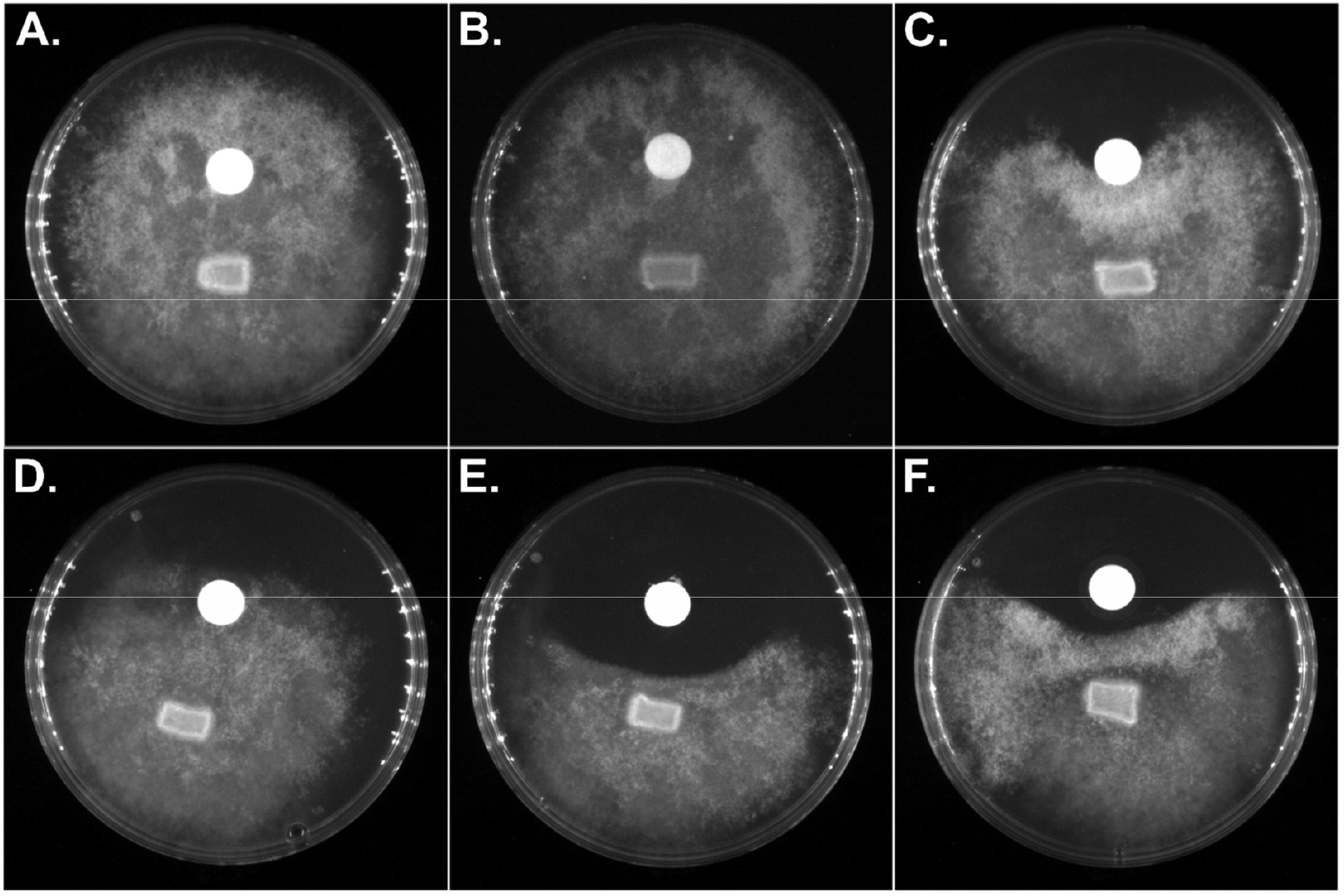
Inhibition assays of fractionation layers from Δ*gacA P. donghuensis* NRC29 complemented with *gacA* carried on pSEVA231 against *A. euteiches*. **A**. Freeze-dried culture. **B**. 80% EtOH precipitation fraction. **C**. 80% EtOH soluble fraction. **D**. Water elution from C18 SPE column. **E**. 50% methanol elution from C18 SPE column. **F**. 100% methanol elution from C18 SPE column.

**Supplementary Table 1.**
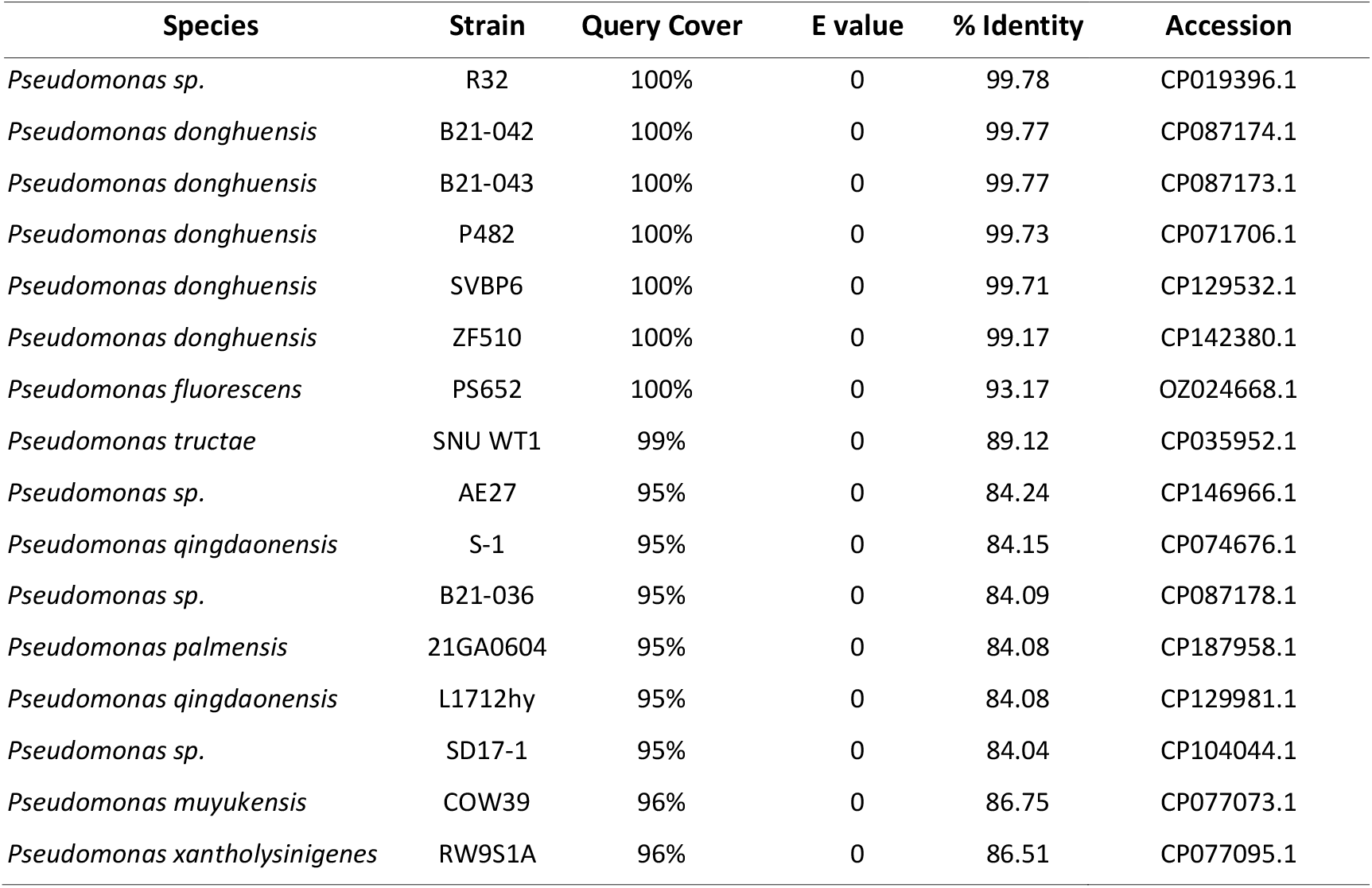
7-hydroxytropolone homologs identified in the core_nt database.

**Supplementary Table 2.**
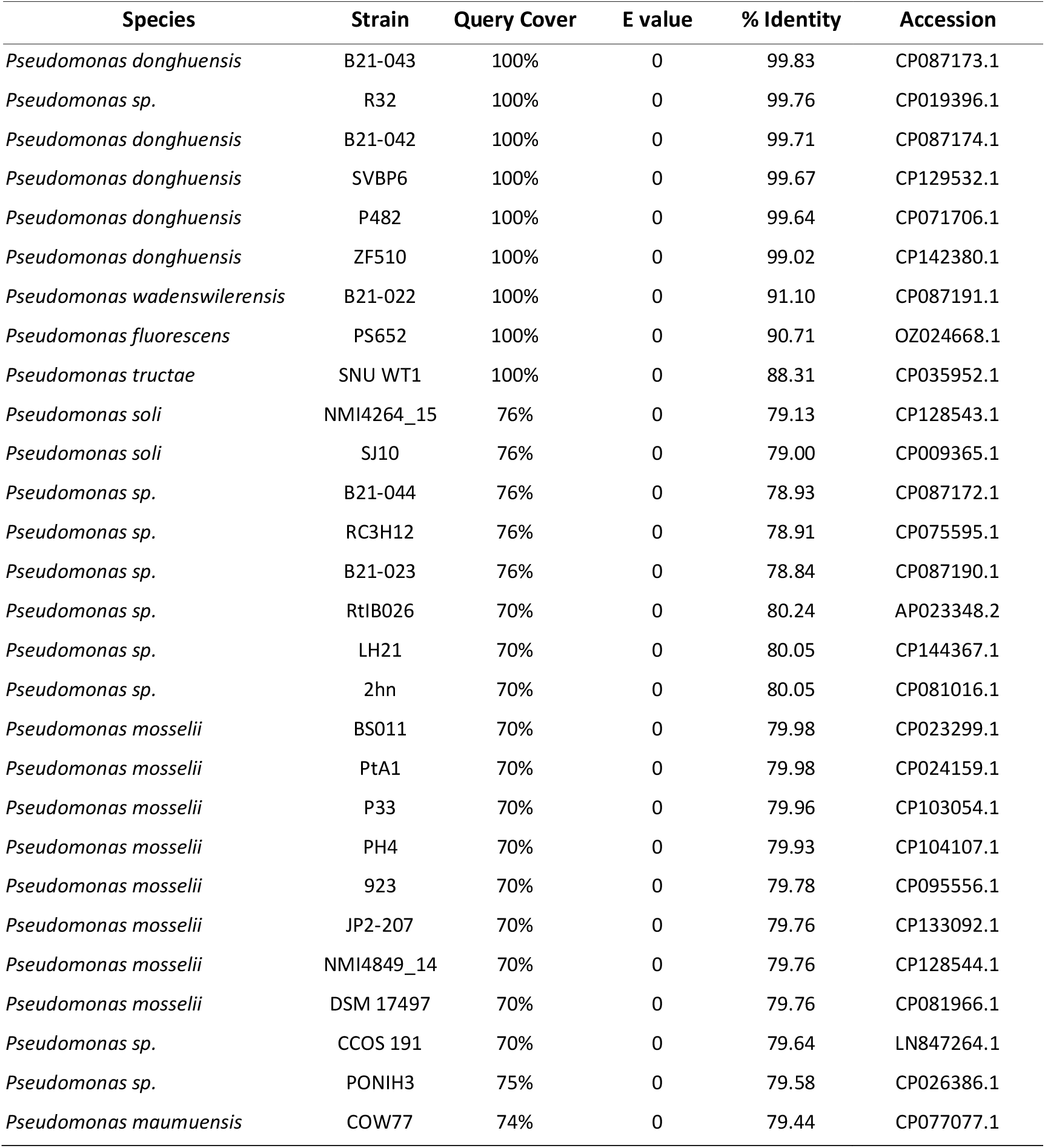
Pseudoiodinine homologs identified in the core_nt database.

